# Cytomegalovirus inhibitors of programmed cell death prevent a contribution of antigen cross-presentation to the priming of antiviral CD8 T cells

**DOI:** 10.1101/2024.04.09.588745

**Authors:** Stefan Ebert, Verena Böhm, Julia K. Büttner, Wolfram Brune, Melanie M. Brinkmann, Rafaela Holtappels, Matthias J. Reddehase, Niels A. Lemmermann

**Affiliations:** Institute for Virology, University Medical Center of the Johannes Gutenberg-University Mainz, Mainz, Germany; Leibniz Institute of Virology (LIV), Hamburg, Germany; Institute of Genetics, Technische Universität Braunschweig, Braunschweig, Germany; Virology and Innate Immunity Research Group, Helmholtz Centre for Infection Research, Braunschweig, Germany; Research Center for Immunotherapy (FZI), University Medical Center of the Johannes Gutenberg-University Mainz, Mainz, Germany; Institute of Virology, Medical Faculty, University of Bonn, Bonn, Germany

## Abstract

CD8 T cells are the predominant effector cells of adaptive immunity in preventing cytomegalovirus (CMV) multiple-organ disease caused by cytopathogenic tissue infection. The mechanism by which CMV-specific, naïve CD8 T cells become primed and clonally expand is of fundamental importance for our understanding of CMV immune control. For CD8 T-cell priming, two pathways have been identified: direct antigen presentation by infected professional antigen-presenting cells (pAPCs) and antigen cross-presentation by uninfected pAPCs that take up antigenic material derived from infected tissue cells. Studies in mouse models using murine CMV (mCMV) and precluding either pathway genetically or experimentally have shown that, in principle, both pathways can congruently generate the mouse MHC/H-2 class-I-determined epitope-specificity spectrum of the CD8 T-cell response. Own recent studies, however, have shown that direct antigen presentation is the canonical pathway when both are accessible. This raised the question of why antigen cross-presentation is ineffective even under conditions of high virus replication thought to provide high amounts of antigenic material for feeding cross-presenting pAPCs. As delivery of antigenic material for cross-presentation is associated with programmed cell death, and as CMVs encode inhibitors of different cell death pathways, we pursued the idea that these inhibitors restrict antigen delivery and thus CD8 T-cell priming by cross-presentation. To test this hypothesis, we compared the CD8 T-cell responses to recombinant mCMVs lacking expression of the apoptosis-inhibiting protein M36 or the necroptosis-inhibiting protein M45 with responses to wild-type mCMV and revertant viruses expressing the respective cell death inhibitors. The data reveal that increased programmed cell death caused by deletion of either M36 or M45 improves CD8 T-cell priming in mice capable of antigen cross-presentation but not in a mutant mouse strain unable to cross-present. These findings strongly support the conclusion that CMV cell death inhibitors restrict the priming of CD8 T cells by antigen cross-presentation.

**Author Summary:** In patients as well as in experimental mouse models, CD8 T cells represent the most potent antiviral effector cells in preventing CMV disease in immunocompromised recipients of hematopoietic cell transplantation. Despite the clinical relevance of mounting a protective response, the mode of CMV antigen presentation to naïve CD8 T cells remained unclear. In principle, naïve CD8 T cells can be sensitized through “direct antigen presentation” on the surface of infected professional antigen-presenting cells (pAPCs) or through “antigen cross-presentation” by uninfected pAPCs that take up antigenic material derived from infected cells following cell death. As CMVs are cytopathogenic, eventually killing their host cells, and as they encode immune evasion proteins interfering with the MHC/HLA class-I pathway of direct antigen presentation in infected cells, it was reasonable to propose a dominant role for antigen cross-presentation. Mouse models precluding either pathway, however, revealed that both can raise an equivalent CD8 T-cell response, and recent work has identified direct antigen presentation as the canonical pathway taken when both are accessible. Here we show that virus-encoded inhibitors of two programmed cell death modalities, apoptosis and necroptosis, prevent an antigen release sufficient for a notable cross-presentation. This answers a long-debated open question in CMV immunology.

## Introduction

Medical interest in cytomegaloviruses (CMVs), members of the β-subfamily of the herpes virus family [1], is based on life-threatening disease that the human cytomegalovirus (hCMV) causes in fetuses, newborn infants, and immunocompromised patients [2–4]. Risk groups in transplantation centers worldwide are recipients of hematopoietic cell transplantation (HCT) [5] and solid organ transplantation (SOT) [6], making hCMV a medically relevant pathogen with an impact on individual health and the public health economy [7,8].

CMVs are host species-specific [9–11] as a result of eons of co-evolution with and adaptation to their specific mammalian hosts, leading to the acquisition of “private genes” not shared between different CMV species [1,12]. As far as analyzed, most of these viral private genes are involved in the interaction between the virus and the immune system. Prominent examples include viral immune evasion genes, which are not conserved but host-specific [13,14]. As revealed by the Bayesian evolutionary tree for the genus *Cytomegalovirus* [1,15,16], CMV species have segregated molecularly from a common ancestor and retained homologous genes involved in the viral replication machinery and in the interaction with cellular signaling pathways conserved between mammalian host species. Examples include viral inhibitors of programmed cell death [17–19]. Importantly, not-shared “private genes” often have analogous functions reflecting the evolutionary principle of “biological convergence”. For instance, hCMV and non-human CMVs, such as murine CMV (mCMV), have independently evolved non-homologous genes interfering by quite different molecular mechanisms with the MHC/HLA class-I pathway of direct antigen presentation to CD8 T cells in the form of peptide-loaded MHC class-I (pMHC-I) complexes at the surface of infected cells [13,14,20].

Based on homologous as well as functionally analogous genes, key features of hCMV pathogenesis and immune control can be reproduced in animal CMV-host pairs. These include the facts that (1) productive primary infection is rapidly cleared in the immunocompetent host by effector mechanisms of innate and adaptive immunity without overt clinical symptoms [13], viral genomes are not eliminated in the course of the termination of productive infection but are maintained in certain cell types in a latent state defined by absence of coordinated productive cycle gene expression, and can be reactivated to productive infection [21,22], and primary infection of an immunocompromised host is productive and cytolytic, leading to intra-tissue virus spread with extensive tissue destruction that eventually can cause multiple organ failure with lethal outcome [4,23,24].

For this reason, animal models can serve to make predictions regarding CMV disease, natural immune control, and therapeutic immune intervention in all questions that cannot be addressed by clinical investigation. Examples are experimental studies on the role of host and viral genes by using specifically tailored transgenic host mutant strains and recombinant viruses with mutations in genes of interest. Non-human primate models are closest to hCMV infection [25], but the mouse model based on infection with mCMV is the most versatile, in particular with regard to altering host genetics [26], and thus the most advanced [27]. Although no model can fully reproduce human disease, predictions made from the mouse model have often proven valid also in human infection, not in all molecular details but in convergently evolved principles [14,15,28]. It was up to the mouse model to provide evidence for the protective function of CD8 T cells in preventing cytopathogenic viral intra-tissue spread and histopathology [29–33]. Specifically, the mouse model revealed that timely reconstitution of high-avidity CD8 T cells is crucial for preventing CMV disease in iatrogenically immunocompromised recipients of experimental HCT [34–37]. By this, the mouse model pioneered clinical immunotherapy of reactivated hCMV infection and disease in HCT recipients/patients by adoptive transfer of virus-specific CD8 T cells [38–41], which is the last resort to fight hCMV strains that have become refractory to antiviral pharmacotherapy [42].

Given the importance of CD8 T cells, it may come as a surprise that the mode by which virus-specific but antigen inexperienced, naïve CD8 T cells are primed and clonally expand to mount a protective response is still a matter of debate. In principle, naïve CD8 can be primed by “direct antigen presentation” [43,44], that is, recognition by their T-cell receptors (TCR) of pMHC-I complexes on the surface of infected professional antigen presenting cells (pAPCs), such as myeloid hematopoietic lineage-derived macrophages and dendritic cells (DCs). Alternatively, priming can be through “antigen cross-presentation” [45,46] by uninfected pAPCs that engulf and process antigenic material derived from infected cells of any cell type following cell death, such as apoptotic bodies or necroptotic extracellular vesicles [47,48]. In the end, this also leads to the recognition of cell surface pMHC-I complexes sensitizing naïve CD8 T cells.

Inspired by the finding that all CMVs encode immune evasion proteins that interfere with the cell surface transport of pMHC-I complexes [14,20], thereby inhibiting direct antigen presentation, several reports provided evidence for cross-presentation by uninfected pAPCs [49–54]. Most convincingly in our view, Snyder and colleagues [51] showed that priming by MHC-I-deficient cells infected with a spread-deficient mCMV mutant, thus excluding first and subsequent rounds of direct antigen presentation, primed the full viral epitope-specificity repertoire and epitope hierarchy known for infection of C57BL/6 (MHC/H-2^b^ haplotype) mice. Notably, we recently found essentially the same CD8 T-cell response in the mutant MHC/H-2^b^ haplotype mouse strain C57BL/6-Unc93b1^3d/3d^ (briefly Unc93b1^3d/3d^) that is known to lack endosomal TLR3, 7, and 9 signaling and is impaired in exogenous antigen processing and thus in antigen cross-presentation [55]. As the conclusion from these studies, the CD8 T-cell response can be primed alternatively but congruently by either mode of antigen presentation when the respective other pathway is made unaccessible experimentally or genetically.

In two recent reports [56,57], we have addressed the question which of the two priming pathways is the natural default pathway when both are accessible. By modulation of direct antigen presentation using recombinant viruses in which the key immune evasion gene m152 of mCMV is either deleted or overexpressed, we showed that the magnitude of the CD8 T-cell response in a draining regional lymph node (RLN) of immunocompetent mice [56] as well as during immune reconstitution after experimental HCT [57] is primarily determined by direct antigen presentation.

This leaves us with the unsolved question of why antigen cross-presentation, although possible in principle, does not make a notable contribution to CD8 T-cell priming after mCMV infection. Priming by cross-presentation requires the feeding of cross-presenting pAPCs, such as CD8^+^CD11c^+^ DCs [58–60], with antigenic material released from infected cells following programmed cell death, such as apoptosis and necroptosis [61–64]. Apoptosis may result from extrinsic signaling initiated by ligation of death receptors and activation of caspase-8 or from intrinsic signaling involving the activation of caspase-9, both leading to effector caspases 3, 6, and 7 [61]. Necroptosis results either from extrinsic death receptor signaling or from intrinsic signaling initiated by double-stranded nucleic acid, specifically by ds Z-RNA in the case of mCMV [65], both leading to the activation of the receptor-interacting protein kinase 3 (RIPK3) and the mixed lineage kinase domain-like (MLKL) protein (for a simplified overview, see Fig 1). As CMVs express evolutionarily conserved inhibitors of programmed cell death [19,66], we studied the CD8 T-cell response in the RLN of mice infected with recombinant mCMVs lacking either the viral inhibitor of caspase-8 activation (vICA) M36, which inhibits apoptosis [67–70], or the viral inhibitor of RIPK3-signaling M45, which inhibits necroptosis [69,71,72]. Our results confirm a dominating basal contribution of direct antigen presentation to the antiviral CD8 T-cell response, but also reveal an enhanced response to viruses in which inhibition of apoptosis or necroptosis is relieved by deletion of M36 and M45, respectively. This additive contribution to the CD8 T-cell-response on top of the response to direct antigen presentation is definitely due to antigen cross-presentation, because it is missing in Unc93b1^3d/3d^ mice that are genetically deficient in antigen cross-presentation.

**Fig 1.**
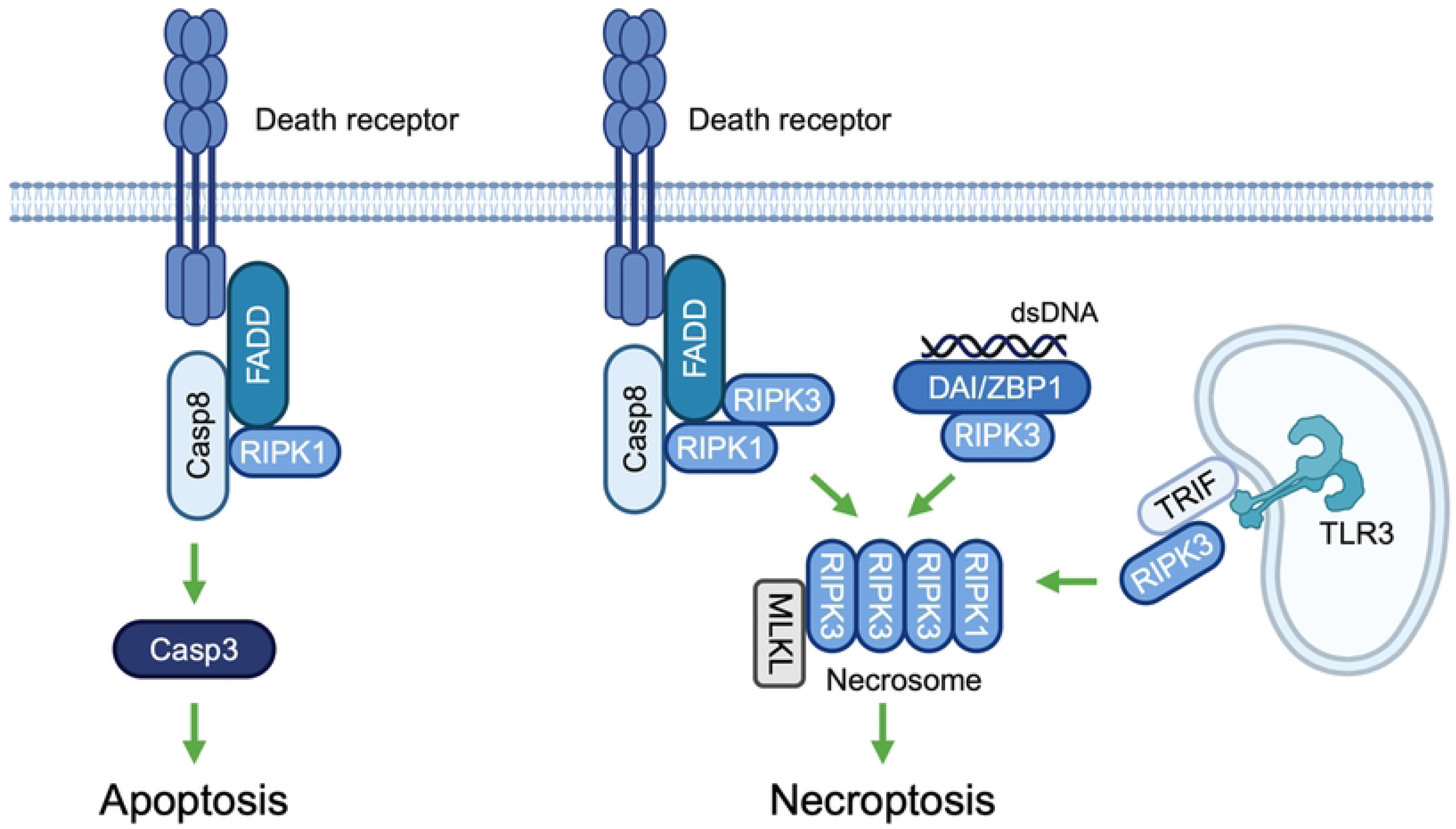
Signaling pathways of programmed cell death: brief introductory overview. (Casp3;8) caspase 3;8, (DAI/ZBP1) DNA-dependent activator of interferon-regulatory factors/Z-DNA-binding protein 1, (FADD) Fas-associated death domain protein, (MLKL) Mixed lineage kinase domain-like protein, (RIPK 1;3) receptor interacting protein kinase 1;3, (TLR3) Toll-like receptor 3, (TRIF) TIR [Toll/interleukin-1 receptor] domain-containing adaptor protein inducing interferon beta. Only the extrinsic signaling pathway is shown for apoptosis.

In conclusion, in normal infection, the viral inhibitors of programmed cell death, namely M36 and M45, prevent a notable contribution of antigen cross-presentation to the antiviral CD8 T-cell response.

## Results

### Deletion of the viral apoptosis inhibitor M36 leads to enhanced CD8 T-cell priming

Viral inhibitor of caspase-8 activation (vICA) M36 of mCMV is the homolog of hCMV vICA UL36 [67,70,73], and UL36 can even replace M36 in its function [74]. Both prevent apoptosis by inhibiting caspase-8 activation (Fig 2A). A possible impact of UL36 on the priming of a human CD8 T-cell response cannot be studied experimentally by infection of volunteers with a UL36 deletion mutant of hCMV. We therefore used the mouse model to study the viral epitope-specific CD8 T-cell response in a draining regional lymph node (RLN), which is the popliteal lymph node in the case of intra-plantar infection. Notably, the anti-apoptotic function of M36 was originally described to operate in macrophages [68,75,76], and macrophages can serve as pAPCs for the priming of a CD8 T-cell response in the RLN after intra-plantar infection with mCMV (S1 Fig).

**Fig 2.**
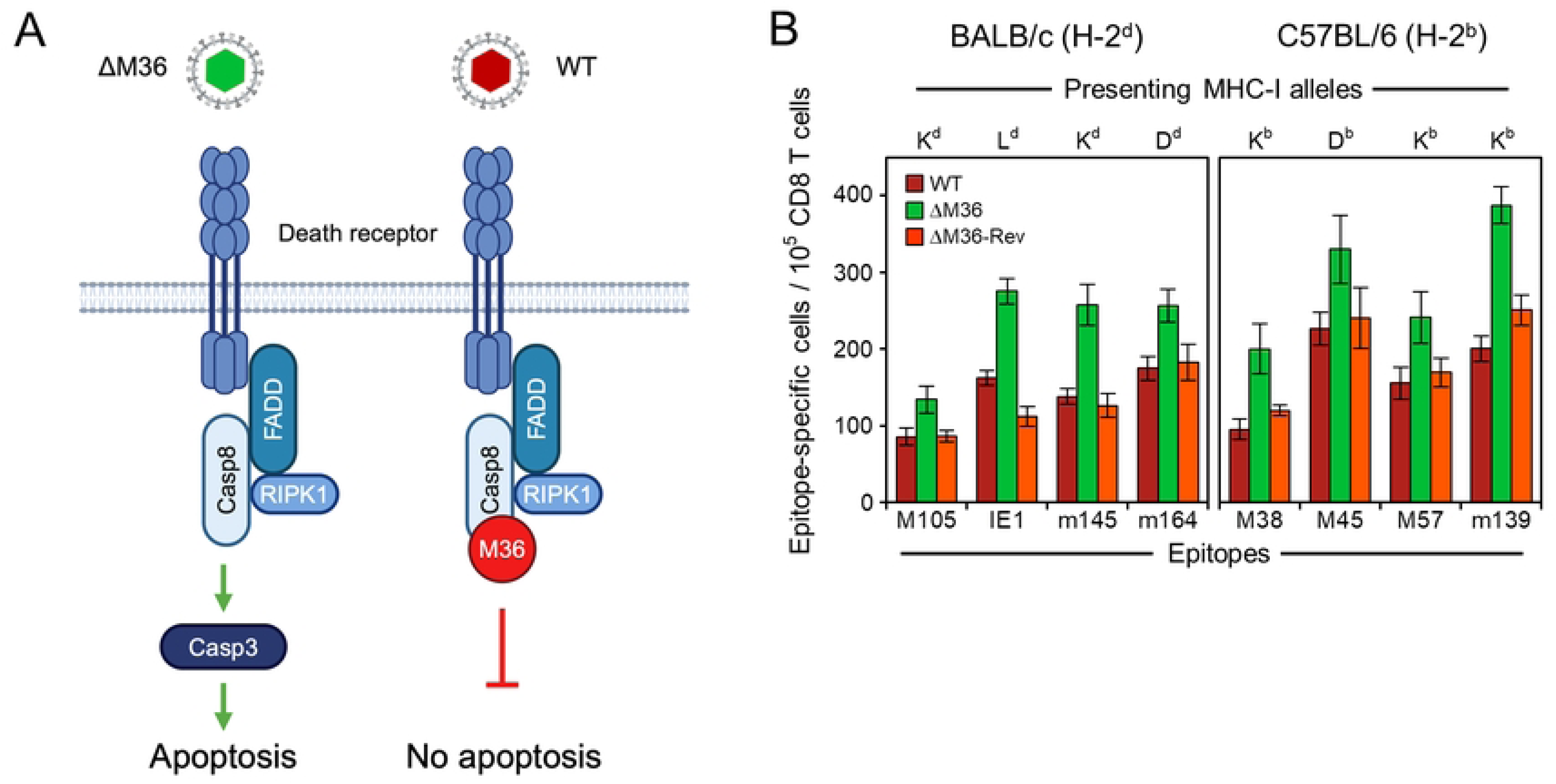
Release from apoptosis blockade by deletion of M36 enhances the antiviral CD8 T-cell response. (**A**) Sketch illustrating the block in the apoptotic signaling pathway by mCMV protein M36. (ΔM36) mCMV-ΔM36; virus symbol colored green to indicate the approval of apoptosis by deletion of M36. (WT) mCMV-WT; virus symbol colored red to indicate the inhibition of apoptosis by M36. (**B**) Viral epitope-specific CD8 T-cell response determined for the RLN, specifically the popliteal lymph node, on day 7 after intra-plantar infection of BALB/c or C57BL/6 mice with viruses mCMV-WT (WT) expressing M36, mCMV-ΔM36 (ΔM36) lacking M36, and the revertant virus mCMV-ΔM36-Rev expressing M36. (Green color) apoptosis can take place. (Red color) apoptosis is blocked. Bars represent most probable numbers, that is, frequencies of responding CD8 T cells measured in ELISpot assays performed for cohorts of 5 mice per infecting virus. Error bars indicate 95% confidence intervals (CI) determined by intercept-free linear regression analysis based on graded numbers of responder cells. Differences are considered significant when the 95% CI do not overlap.

We compared the CD8 T-cell responses to wild-type (WT) mCMV (mCMV-WT), the M36 gene deletion mutant mCMV-ΔM36, and the corresponding revertant virus mCMV-ΔM36-Rev in the RLN of BALB/c (MHC haplotype H-2^d^) mice on day 7 after intra-plantar infection (Fig 2B, left panel). It must be noted that the response magnitude reflects the initial event of the sensitization of naïve CD8 T cells as well as subsequent clonal expansion. For four representative epitopes of mCMV presented as antigenic peptides by MHC class-I (MHC-I) molecules K^d^ (epitopes M105 and m145), L^d^ (epitope IE1), and D^d^ (epitope m164), the response to infection with mCMV-ΔM36 was consistently higher than the response to WT virus. This enhanced response was functionally reverted to WT virus level after infection with the M36-restored virus mCMV-ΔM36-Rev.

In an independent second experiment, the CD8 T-cell responses in the RLN to mutant and revertant virus were compared in the time course after intra-plantar infection of BALB/c mice (S2 Fig). The data essentially reproduced and thus confirmed the results shown for day 7 (Fig 2B) and added the information that minor differences in the initial priming event gain statistical significance over time of clonal expansion, that is, after multiple rounds of proliferation.

The findings were then reproduced in an analogous experimental setting by infection of C57BL/6 (MHC haplotype H-2^b^) mice (Fig 2B, right panel). For four representative epitopes of mCMV presented as antigenic peptides by MHC-I molecules K^b^ (epitopes M38, M57, and m139), and D^b^ (epitope M45), the response to infection with mCMV-ΔM36 was consistently higher than the response to WT virus. Again, this enhanced response was functionally reverted to WT virus level after infection with mCMV-ΔM36-Rev.

In conclusion from this set of data, apoptosis of infected cells enabled by deletion of vICA M36 enhances the CD8 T-cell response in an RLN for all viral epitopes tested for two commonly used mouse inbred strains and independent of the MHC-I molecule that presents the respective antigenic peptide.

### Virus attenuation by deletion of M36 is not caused by enhanced immune control but reflects premature loss of virus-producing cells due to apoptosis

During evolution, CMVs have acquired inhibitors of programmed cell death to avoid premature death of infected cells before progeny virions are assembled and released for cell-to-cell spread. Accordingly, deletion of these inhibitors should lead to viral attenuation. We tested this prediction for the example of M36 (Fig 3). We quantitated spliced immediate-early 1 (IE1) transcripts, which are proportional to the number of infected cells, as a surrogate for the degree of infection. The more direct parameter for viral replication, the copy number of viral DNA, cannot be used in this particular case, because the infection dose of 10^5^ infectious units (plaque-forming units, PFU) equals ∼5 x 10^7^ genomic viral DNA molecules (genome-to-PFU ratio of 500) [77,78] that obscure the detection of *de novo* synthesized viral DNA, especially at early times after infection.

**Fig 3.**
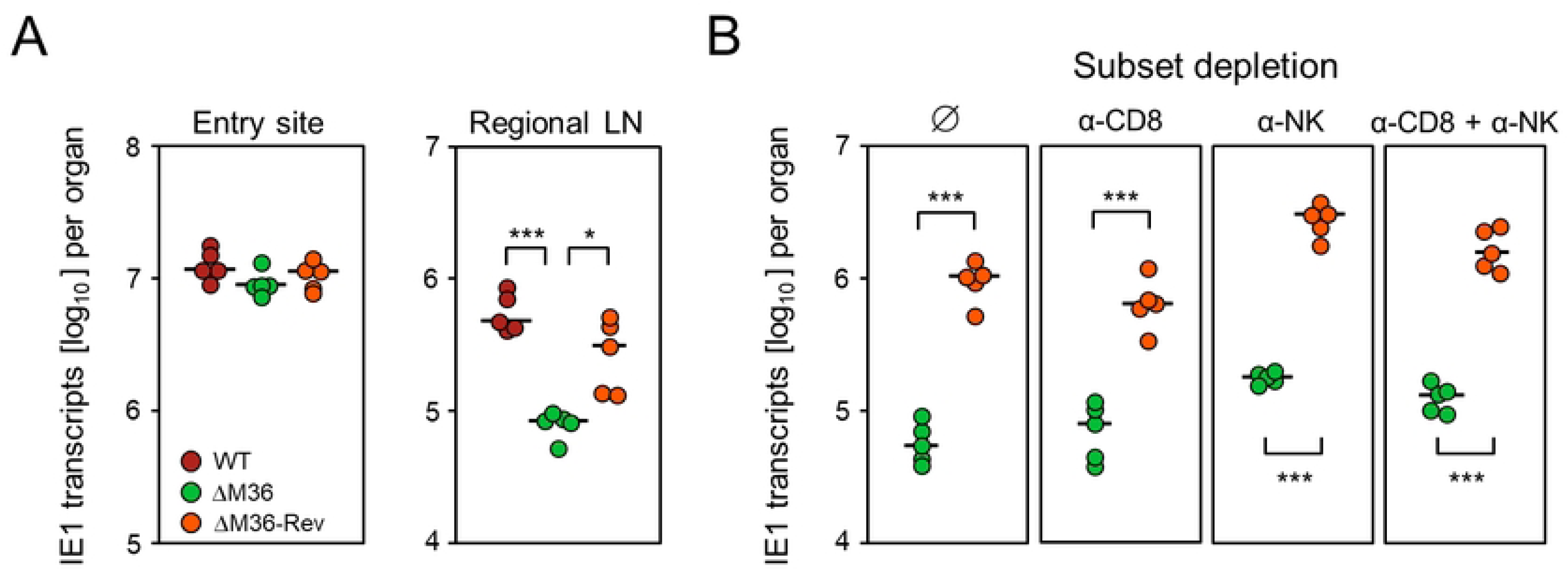
Release from apoptosis blockade by deletion of M36 leads to virus attenuation in the RLN but not locally at the viral entry site. (**A**) Level of infection determined by quantitation of IE1 transcripts in plantar (footpad) tissue (left panel) and in the draining RLN, the popliteal lymph node (right panel), at 48 hours (day 2) after intra-plantar infection (day 0). (**B**) Level of infection determined by quantitation of IE1 transcripts in the RLN at 72 hours (day 3) after intra-plantar infection (day 0). At 24 hours before infection (day -1), mice were depleted of CD8 T cells (panel α-CD8) or NK cells (panel α-NK) or both (panel α-CD8 + α-NK). (Panel ∅), control group with no depletion. Infecting viruses are indicated and color-coded as in Fig 2B. Filled circles represent data from 5 mice per experimental group (n = 5) tested individually. Median values are marked. Levels of significance: P-values (*) <0.05 and (***) <0.001.

On day 2 after intra-plantar infection, a time allowing one round of viral replication in plantar tissue [79], local IE1 gene expression was identical for WT virus as well as ΔM36 mutant and revertant viruses (Fig 3A, left panel). This indicates an equivalent viral capacity to infect cells at the entry site and suggests resistance to apoptosis of infected connective tissue cells. In contrast, at the same time, the M36 deletion mutant was significantly growth-attenuated compared to WT virus and the revertant virus in the popliteal lymph node, that is, the draining RLN (Fig 3A, right panel). Notably, previous work has shown that mCMV-ΔM36 fails to disseminate to the liver after intra-plantar infection, unless inhibition of apoptosis in cells infected with the mutant virus is restored by functional M36 trans-complementation upon co-infection with WT virus after ipsilateral but not after contralateral intra-plantar application of both viruses [80].

In accordance with the function of M36, attenuation of the deletion mutant likely reflects reduced virus production and spread due to premature death of infected cells by apoptosis. We also considered the alternative explanation of an enhanced immune control, selectively of the mutant virus, by viral antigen cross-presentation to CD8 T cells. This idea was refuted, however, by depletion of CD8 T cells, natural killer (NK) cells, or both (Fig 3B). While the depletion of CD8 T cells had no notable effect on early viral replication (Fig 3B, first two panels from the left), the depletion of NK cells indicated some early control by innate immunity (Fig 3B, first and third panel from the left). The difference between mutant and revertant virus, however, was not abolished by any immune cell subset depletion (Fig 3B, all panels).

In conclusion from this set of data, attenuation of mCMV-ΔM36 in the RLN is not caused by an enhanced immune control but is consistent with apoptosis of infected cells preventing virus release and spread.

### Alternative inhibition of apoptosis by blocking FADD signaling reverts the effect of M36 deletion

We considered the theoretical possibility that deletion of M36 might improve the CD8 T-cell response by a mechanism unrelated to enhanced apoptosis. To test this, inhibition of apoptosis was restored in absence of M36 by viral expression of a dominant-negative mutation of FADD (FADD^DN^), which competes with cellular FADD in cells infected with recombinant virus mCMV-ΔM36.FADD^DN^ [75] (Fig 4A).

**Fig 4.**
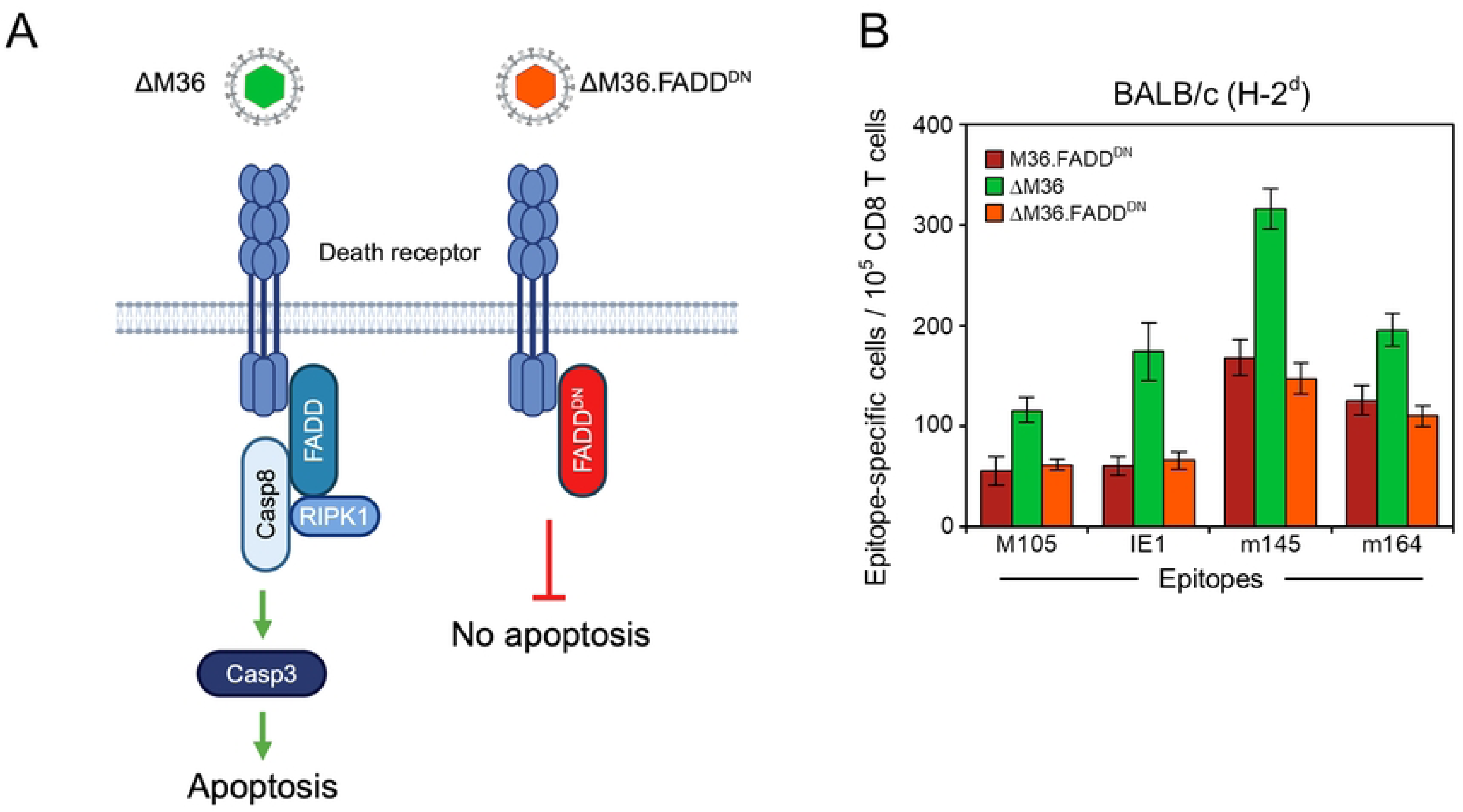
Interruption of apoptotic signaling by virally-encoded dominant-negative FADD functionally reverts the enhancing effect of M36 deletion on the antiviral CD8 T-cell response. (**A**) Sketch illustrating the block in the apoptotic signaling pathway by replacement of cellular FADD with virally-encoded FADD^DN^. (ΔM36) mCMV-ΔM36; virus symbol colored green to indicate the approval of apoptosis by deletion of M36. (ΔM36.FADD^DN^) mCMV-ΔM36.FADD^DN^; virus symbol colored red to indicate the interruption of signaling by FADD^DN^. (**B**) Viral epitope-specific CD8 T-cell response determined for the RLN, specifically the popliteal lymph node, on day 7 after intra-plantar infection of BALB/c mice (cohorts of 5 mice) with the viruses indicated in the internal legend. (Green color) apoptosis can take place. (Red color) apoptosis is blocked. For further information, see the legend of Fig 2B.

Analogous to the experiment shown in Fig 2B, the CD8 T-cell response was tested in the RLN of BALB/c mice on day 7 after intra-plantar infection (Fig 4B). Compared to a control virus expressing M36 and FADD^DN^, the CD8 T-cell response was consistently enhanced by infection with mCMV-ΔM36 for the panel of viral epitopes tested, and this was reversed by expression of FADD^DN^ after infection with virus mCMV-ΔM36.FADD^DN^.

From this, we conclude that the mechanism by which deletion of M36 increases the CD8 T-cell response is indeed related to enhanced apoptosis.

### Enhancement of apoptosis by deletion of M36 leads to a higher and broader antiviral CD8 T-cell response

So far, we tested the impact of M36 deletion on the CD8 T-cell response in the RLN with a selected panel of mCMV epitopes for which the corresponding antigenic peptides have been identified in their amino acid sequence (for an overview, see [81]). To get an impression of the overall antiviral CD8 T-cell response, we used an mCMV genome-wide open reading frame (ORF) library of expression plasmids [82] using responder CD8 T cells derived from the spleen on day 7 after intra-plantar infection with viruses preventing or allowing apoptosis (Fig 5, for the assay setup, see [83]). Obviously, compared to viruses preventing apoptosis by expression of M36 or FADD^DN^, the overall antiviral CD8 T-cell response was higher and broader with the vICA M36 deletion virus mCMV-ΔM36.

**Fig 5.**
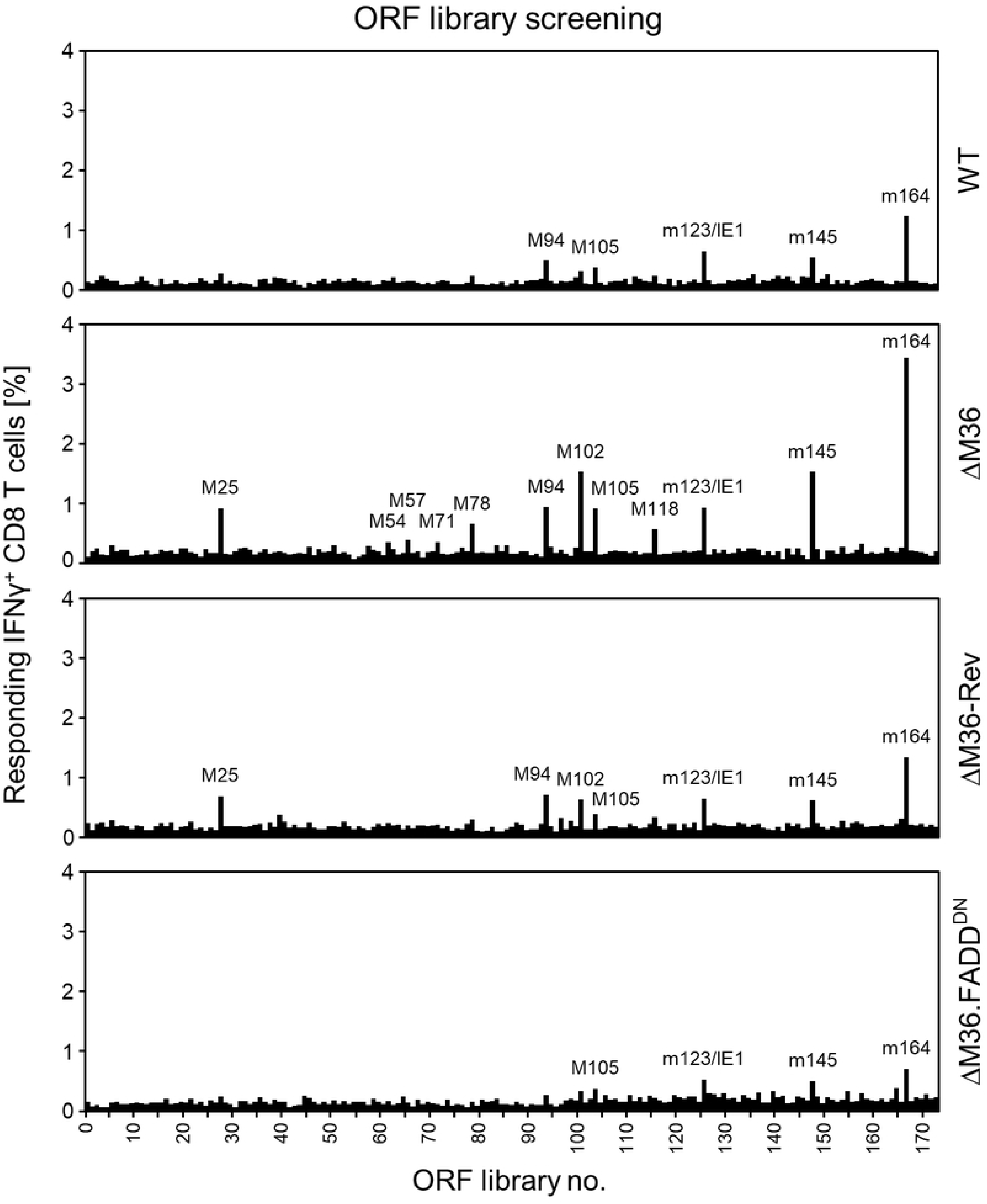
Viral genome-wide screening of the viral antigen-specific CD8 T-cell response in dependence on apoptosis. A viral genome-spanning ORF library of expression plasmids was used to test the overall CD8 T-cell response determined for the spleen on day 7 after intra-plantar infection of BALB/c mice (cohorts of 7 mice) with the viruses indicated. Signals represent frequencies of CD8 T cells activated to express intracellular IFNγ after stimulation with antigenic transfectants. For more prominent signals, the corresponding ORF is named.

### Enhancement of necroptosis by inactivation of M45 leads to a generally higher and broader antiviral CD8 T-cell response

We next addressed the question if the impact of programmed cell death on the CD8 T-cell response applies selectively to apoptosis or also to necroptosis (recall Fig 1). The viral inhibitor of RIPK3-signaling M45 of mCMV prevents the formation of the necroptosis signaling complex, also known as necrosome [61], particularly in macrophages (Fig 6A) [69]. It is important to note that necroptosis after deletion of M45 comes into charge only when apoptosis is blocked by M36 [19,61,66,71].

**Fig 6.**
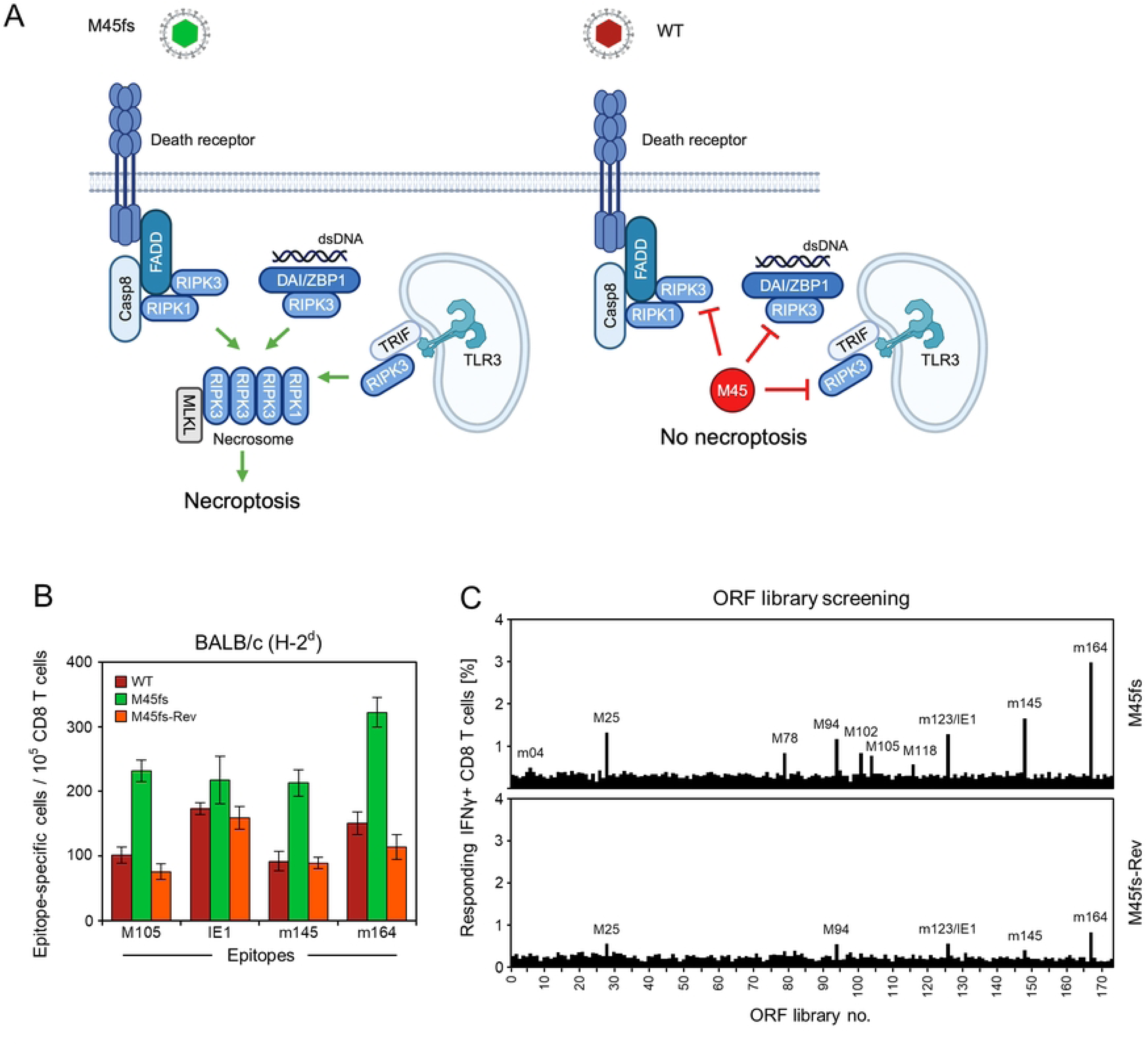
Release from necroptosis blockade by functional deletion of M45 enhances the antiviral CD8 T-cell response. (**A**) Sketch illustrating the block in the necroptotic signaling pathway by mCMV protein M45. (M45fs) mCMV-M45fs; virus symbol colored green to indicate the approval of necroptosis by functional deletion of M45. (WT) mCMV-WT; virus symbol colored red to indicate the inhibition of necroptosis by functional M45. (**B**) Viral epitope-specific CD8 T-cell response determined for the RLN, specifically the popliteal lymph node, on day 7 after intra-plantar infection of BALB/c mice (cohorts of 5 mice) with viruses mCMV-WT (WT) expressing functional M45, mCMV-M45fs (M45fs) lacking functional M45, and the revertant virus mCMV-M45fs-Rev (M45fs-Rev) expressing functional M45. (Green color) necroptosis can take place. (Red color) necroptosis is blocked. For further information, see the legend of Fig 2B. (**C**) ORF library screening of the overall CD8 T-cell response determined for the spleen on day 7 after intra-plantar infection of BALB/c mice (cohorts of 7 mice) with the viruses indicated. For further information see the legend of Fig 5.

We compared the CD8 T-cell responses in the RLN on day 7 after intra-plantar infection with the necroptosis-inducing virus mCMV-M45fs, in which M45 is inactivated by replacing a transposon insertion with an M45 gene frame-shift mutation [84], and the necroptosis-inhibiting viruses mCMV-WT and mCMV-M45fs-Rev (Fig 6B). As it was the case for the apoptosis inhibitor M36 (recall Fig 2B), necroptosis in absence of a functional M45 led to an enhanced CD8 T-cell response compared to mCMV-WT for all epitopes tested, and was reversed for mCMV-M45fs-Rev expressing M45. This analogy between inhibition of apoptosis and inhibition of necroptosis also extends to virus attenuation after deletion of the respective cell death inhibitor (S3 Fig, compared to Fig 3A).

To get an impression of the overall antiviral CD8 T-cell response, we again used the genome-wide ORF-library of expression plasmids (recall Fig 5) to screen the response by CD8 T cells derived from the spleen on day 7 after intra-plantar infection (Fig 6C). Obviously, the response was higher and broader after infection with mCMV-M45fs compared to mCMV-M45fs-Rev.

### Deletion of programmed cell death inhibitors enhances the CD8 T-cell response by antigen cross-presentation

All data have consistently shown that programmed cell death results in an enhanced antiviral CD8 T-cell response. It is well-established that apoptotic bodies and necroptotic extracellular vesicles can be taken up efficiently by mature, uninfected pAPCs for antigen cross-presentation to prime naïve CD8 T cells [85–87]. It was therefore reasonable to assume that the enhanced CD8 T-cell response observed after deletion of M36 or M45 was caused by antigen cross-presentation.

To replace this educated guess with firm evidence, we tested all deletion mutants and corresponding revertants in mouse strain Unc93b1^3d/3d^ genetically deficient in antigen cross-presentation [55].

The strong CD8 T-cell response observed in these mice (Fig 7A-C) is thus based entirely on direct antigen presentation and proves that direct presentation remains the predominant mode of CD8 T-cell priming also when apoptosis or necroptosis can take place. The most important conclusion of these experiments is that apoptosis or necroptosis do not enhance the CD8 T-cell response when antigen cross-presentation is precluded. Conversely, the results indicate that the enhancement of the response observed in mice that are genetically competent in cross-presentation indeed resulted from cross-presentation as an add-on to an underlying direct antigen presentation.

**Fig 7.**
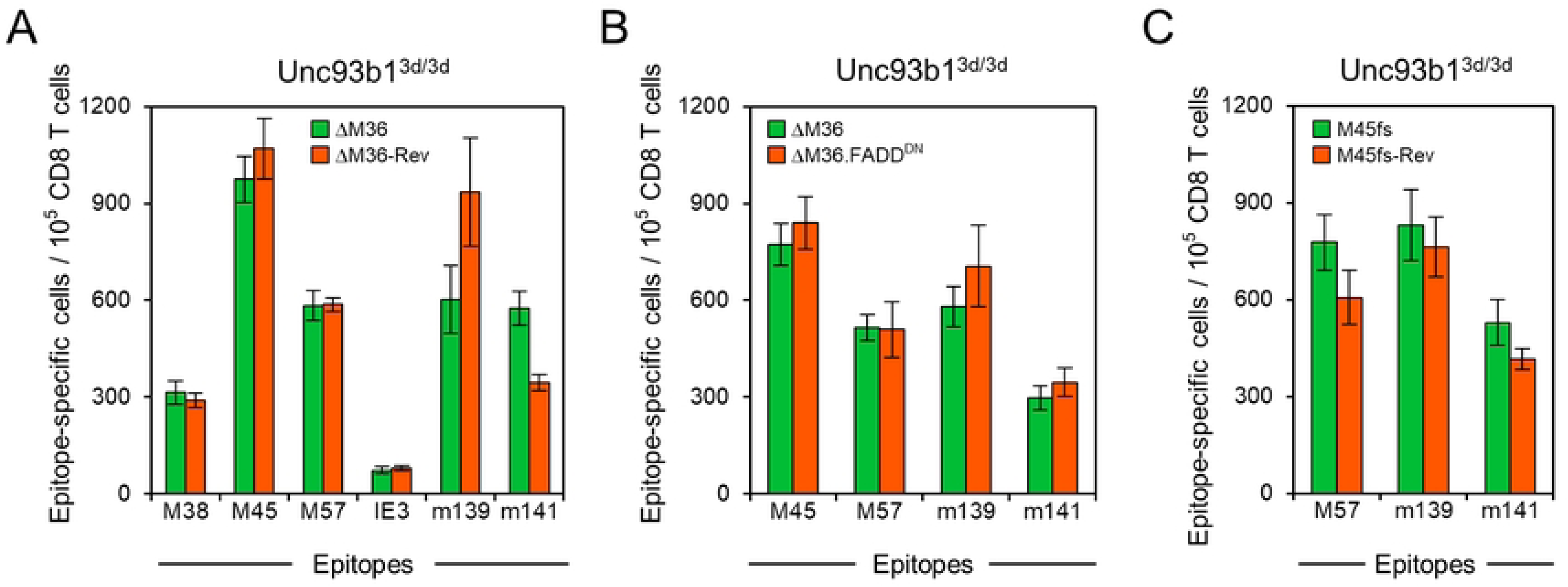
Missing contribution of programmed cell death to the CD8 T-cell response in C57BL/6-Unc93b1^3d/3d^ mice genetically unable to cross-present. (**A**) Blockade of apoptosis by M36. (B) Blockade of apoptosis by FADD^DN^. (**C**) Blockade of necroptosis by M45. (**A-C**) Viral epitope-specific CD8 T-cell response determined for the RLN, specifically the popliteal lymph node, on day 7 after intra-plantar infection of C57BL/6-Unc93b1^3d/3d^ mice (cohorts of 4 mice) with the viruses indicated. Note that A-C represent independent experiments. For further information, see the legend of Fig 2B.

## Discussion

The way how naïve CD8 T cells become primed to mount a protective antiviral response has long been a matter of debate in CMV immunology. Evidence was presented supporting direct antigen presentation by infected pAPCs as well as antigen cross-presentation by uninfected pAPCs that engulf antigenic material derived from infected cells undergoing programmed cell death and releasing apoptotic bodies or necroptotic extracellular vesicles. However, proponents of one mechanism or the other have used sophisticated experimental protocols or mutant mouse strains to rule out the respective alternative mechanism. Specifically, experimental prevention of direct antigen presentation showed priming by cross-presentation [51,52,88], and prevention of cross-presentation showed priming by direct presentation [56,89]. The bottom line was that, in principle, both mechanisms can generate antiviral CD8 effector T cells recognizing essentially the same viral epitopes in the same immunodominance hierarchy [51,56]. However, which type of priming prevails in normal mice competent for both direct and cross-presentation remained an open question.

By using the new strategy to modulate direct antigen presentation by mCMV mutants where the key immune evasion gene m152 was either deleted or overexpressed, we recently showed that the CD8 T-cell response directly reflects the level of antigen presentation in the form of pMHC-I complexes on the surface of infected cells [56,57]. This applied to priming of naïve CD8 T cells in lymphoid tissue, specifically in the RLN of immunocompetent mice [56], but also to priming in the medically relevant situation after HCT when CD8 T cells become reconstituted in hematoablated HCT recipients only gradually over time [57]. However, it remained the question why antigen cross-presentation does not seem to play a significant role under physiological conditions.

Here we addressed this open question by studying mCMV mutants with deletion of the viral cell death inhibitors M36 or M45. For mouse strains genetically capable of antigen cross-presentation, such as BALB/c and C57BL/6, our data show an enhanced CD8 T-cell response to mutant viruses that induce apoptosis or necroptosis. This response increment is lost in Unc93b1^3d/3d^ mice [55] genetically deficient in antigen cross-presentation (Fig 7.) Thus, viral inhibition of programmed cell death explains the lack of antigen cross-presentation in normal mice. As a by-product, direct antigen presentation in Unc93b1^3d/3d^ mice was not negatively affected by programmed cell death (Fig 7), although one would expect a reduction in the number of antigen-presenting pAPCs. To explain this unexpected phenomenon, we can currently only speculate that priming by direct antigen presentation might be completed before the infected pAPCs are lost by programmed cell death. Another topic of discussion is the question of why the “late death” of productively infected cells by cytolysis as the endpoint of the cytopathogenicity of CMVs does not seem to support antigen cross-presentation. A reasonable possibility might be that cell debris resulting from cytolytic cell death, unlike apoptotic bodies or necroptotic extracellular vesicles, lacks signals or ligands that induce uptake by cross-presenting pAPCs [90,91].

Programmed death of infected cells initiated by the expression of death receptor ligands is an antiviral defense mechanism of the host to prevent viral spread. Thus, from the perspective of viral evolution, acquisition of genes that inhibit apoptosis and necroptosis enables the virus to prevent premature cell death, and thereby allows completion of the viral productive cycle and virus exit before the infected cells eventually die a “late death” from the cytolytic infection [19].

Previous work on the function of M36 and M45 emphasized their role in “viral fitness” as a pathogenicity factor and determinant of cell tropism [84] by showing “viral attenuation” in case of their deletion [68,69,72,75,84,92–96]. Interestingly, in the case of M36 deletion, attenuation is also associated with greater susceptibility to effector CD8 T-cell control, based on a cross-talk between apoptotic signaling and perforin/granzyme-dependent cytotoxicity [70]. In accordance with these studies, we document an early viral attenuation in the RLN by deletion of M36 (Fig 3A) or M45 (S3 Fig), although at least this early effect was not mediated by an antiviral immune control (Fig 3B). As a new aspect, our data add the information that cell death inhibitors contribute to viral spread and pathogenesis not only by preventing premature cell death to allow completion of the viral replication cycle but, in addition, also by preventing antigen cross-presentation leading to a suboptimal CD8 T-cell response.

We were initially wondering why attenuation by deletion of M36 or M45 is not seen at the viral entry site, which is the footpad tissue in the case of intra-plantar infection (Fig 3A, S3 Fig). However, there exists an explanation. Virions administered into the interstitial fluid do not need to spread from infected footpad connective tissue to the RLN after a first round of local infection, but can directly reach the RLN within minutes via afferent lymphatic vessels [97,98]. Therefore, the observed difference in virus attenuation most likely results from different susceptibility to programmed cell death of the cell types infected in footpad tissue and RLN.

Murine fibroblasts and endothelial cells were reported to undergo apoptosis upon mCMV-ΔM36 infection only after conditioning by tumor necrosis factor α (TNFα) [99]. Our functional data on the lack of an M36 deletion phenotype in the footpad (Fig. 3A) suggest that infected connective tissue fibrocytes or infected endothelial cells in the footpad vasculature are not susceptible to ΔM36-induced apoptosis. In this context is important to recall that M36 and M45 were originally both found to operate in infected macrophages [68,76,96], and M36 was found to block apoptosis also in DCs [99]. After intravenous infection of immunocompromised BALB/c mice, mCMV-ΔM36 was found to replicate in liver tissue, and apoptosis was detected *in situ* by staining of active caspase-3 in Kupffer cells, which are the liver macrophages, but also in hepatocytes, which are the liver parenchymal cells [75]. Importantly, macrophages and DCs are natural sources of TNFα that trigger apoptosis in an autocrine fashion after infection with mCMV-ΔM36 [99]. In this context, it is important to note that virions reaching the RLN via the lymphatics are deposited in the RLN subcapsular sinus (SCS) lined with SCS macrophages and DCs ([98] and references therein). Moreover, infected macrophages are known determinants of viral pathogenesis [100], and in a related model, mCMV was found to infect CD169^+^ SCS macrophages in mediastinal lymph nodes after systemic infection [101]. From all this evidence we propose that attenuation of virus mCMV-ΔM36 in the RLN is caused by apoptosis of infected SCS macrophages and, likely, also SCS DCs. Importantly, the same rules apply to necroptosis by deletion of M45 under the condition that M36 is expressed to prevent apoptosis (S3 Fig).

Our key result of CD8 T-cell priming by antigen cross-presentation, dependent upon either apoptosis or necroptosis after deletion of programmed cell death inhibitors M36 or M45, respectively, is based on consistent data throughout. Uninfected pAPCs that take up apoptotic bodies or necroptotic extracellular vesicles, respectively, in the RLN following intra-plantar infection can be cross-presentation-competent SCS DCs, for instance, CD8^+^CD11c^+^ DCs [58,59], but also SCS macrophages identified more recently as being also capable to cross-present [102].

One should always keep in mind that studying viral mutants does not aim to understand the biology of the mutants but to learn about the evolutionary successful WT virus. While our data have shown that mCMV-specific CD8 T cells can be primed by antigen cross-presentation, provided that apoptosis or necroptosis are facilitated by deletion of the viral inhibitors, the relevant message for understanding the CD8 T-cell response to WT virus is the reverse. CD8 T-cell priming to mCMV-WT occurs predominantly, if not exclusively, by direct antigen presentation, because viral cell death inhibitors prevent a significant contribution of antigen cross-presentation. While antigen cross-presentation would have been in the host’s interest, ensuring viral replicative capacity by simultaneously blocking premature cell death and antigen cross-presentation was likely a driver of selection in CMV evolution.

Our results thus provide an answer to a long-disputed fundamental question in CMV immunology.

## Materials and methods

### Ethics statement

Animal experiments were performed in accordance with the national animal protection law (Tierschutzgesetz (TierSchG)), animal experiment regulations (Tierschutz-Versuchstierverordnung (TierSchVersV)), and the recommendations of the Federation of European Laboratory Animal Science Association (FELASA). The experiments were approved by the ethics committee of the Landesuntersuchungsamt Rheinland-Pfalz, permission numbers 17-07-04/051-62 and 177-07/G09-1-004.

### Cells, viruses, and mice

P815 (No. TIB-64, haplotype H-2^d^) and EL4 (No. TIB-39, haplotype H-2^b^) cells were obtained from the American Tissue Culture Collection (ATCC) and cultivated in RPMI supplemented with 5% fetal calf serum (FCS) and antibiotics, or in DMEM with 10% FCS and antibiotics, respectively. Primary murine embryo fibroblasts (MEF) were cultivated in MEM supplemented with 10% FCS and antibiotics.

Virus derived from BAC plasmid pSM3fr [103] was used as “wild-type (WT)” virus, mCMV-WT. Recombinant viruses mCMV-ΔM36 and mCMV-ΔM36 [68], as well as mCMV-M45.BamX and mCMV-M45.BamX-Rev [84] have been described previously. mCMV-M45.BamX carries a frameshift mutation in M45 [84] and is, for simplicity, renamed mCMV-M45fs in this study. Accordingly, the corresponding revertant virus is renamed mCMV-M45fs-Rev. Recombinant viruses mCMV-ΔM36-FADD^DN^ and mCMV-WT-FADD^DN^ [75], here renamed mCMV-ΔM36.FADD^DN^ and mCMV-M36.FADD^DN^, were used as functional revertants of mCMV-ΔM36.

BALB/ and C57BL/6 mice were bred and housed under specified-pathogen-free (SPF) conditions in the Translational Animal Research Center (TARC) at the University Medical Center of the Johannes Gutenberg-University Mainz, Germany. C57BL/6-Unc93b1^3d/3d^ (briefly Unc93b1^3d/3d^ [55]) mice were bred and housed at the central animal facility of HZI Braunschweig, Germany.

### Depletion of leukocyte subsets *in vivo*

Depletion of NK cells or CD8 T cells was performed 24 h prior to infection by i.v. injection of 25µl rabbit antiserum directed against asialo-GM1 (catalog no. 986-100001; Wako Chemicals, Osaka, Japan) or of 1mg purified antibody directed against CD8 (clone YTS169.4), respectively [104]. Macrophage depletion was performed by intra-plantar injection of 50µl clodronate liposomes (catalog no. C-005 ClodronateLiposomes.com, Haarlem, The Netherlands) or control liposomes (catalog no. P-005) 3 days prior to infection.

### Quantification of viral transcription in infected tissue

Spliced IE1 (m123/ie1) transcripts were quantitated from total RNA extracted from infected lymph nodes [104] or infected footpads using the RNeasy Lipid Tissue Mini Kit (catalog no. 74804 QIAGEN). 500 ng RNA was used as template for RT-qPCR performed with splice product-specific primers and probes. Absolute quantification of IE1 using *in vitro* transcripts as standard has been described previously [79]. Note that transcription is the more sensitive measure of infection compared to the classical PFU assay and, unlike quantitation of viral genomes, it is not obscured by input virion DNA.

### Peptides

Custom peptide synthesis to a purity of > 80% was performed by JPT Peptide Technologies (Berlin, Germany). Synthetic peptides representing antigenic peptides in mouse haplotype H-2^b^ were M38 (SSPPMFRVP), M45 (HGIRNASFI), M57 (SCLEFWQRV), M122/IE3 (RALEYKNL), m139 (TVYGFCLL), and m141 (VIDAFSRL) [82,105]. Those for mouse haplotype H-2^d^ were M105 (TYWPVVSDI), M123/IE1 (YPHFMPTNL), m145 (CYYASRTKL), and m164 (AGPPRYSRI) [81]. The synthetic peptides were used for exogenous loading of stimulator cells in the ELISpot assay.

### ELISpot assay

An interferon gamma (IFNγ) enzyme-linked immunospot (ELISpot) assay was performed for quantification of IFNγ-secreting CD8 T cells after sensitization by peptide-loaded stimulator cells. Frequencies of mCMV-specific CD8 T cells were determined by incubation of graded numbers of immunomagnetically-purified CD8 T cells, derived from the RLN, which is the popliteal lymph node in the case of intra-plantar infection, with stimulator cells (P815/H-2^d^ or EL4/H-2^b^, as it applied) that were exogenously loaded with synthetic peptides at a saturating concentration of 10^-7^M [104]. Spots were counted automatically based on standardized criteria using Immunospot S4 Pro Analyzer (CTL, Shaker Heights, OH, USA) and CTL-Immunospot software V5.1.36. Frequencies (most probable numbers) of IFNγ-secreting cells and the corresponding 95% confidence intervals were calculated by intercept-free linear regression analysis using Mathematica, version 8.0.4 [104,106].

### Genome-wide ORF library screening by intracellular cytokine assay

An mCMV ORF library of expression plasmids spanning the entire mCMV genome [82] was used for ORF-specific stimulation of *ex vivo* isolated CD8 T cells with transfected SV-40 fibroblasts, followed by cytofluorometric (CFM) detection of intracellular IFNγ. The principle and the methods of the ORF library screening have been published previously [82,107].

### Statistical analysis

To evaluate statistical significance of differences between two independent sets of data, the unpaired t-test with Welch’s correction of unequal variances was used. Differences were considered statistically significant for P-values (*) <0.05, (**) <0.01, and (***) <0.001. Calculations were performed with Graph Pad Prism 10.1 (Graph Pad Software, San Diego, CA). In cohort analyses of viral epitope-specific CD8 T cells by the ELISpot assay, differences are considered statistically significant if 95% confidence intervals do not overlap.

## Acknowledgments

The authors thank Angelique Renzaho (Institute for Virology, University Medical Center Mainz) for superb technical assistance and Bernado Franklin (Institute of Innate Immunity, University of Bonn) for helpful discussions. Sketches were designed with BioRender.com.

## Author Contributions

**Conceptualization:** Rafaela Holtappels, Matthias J. Reddehase. Niels A. Lemmermann

**Data Curation:** Stefan Ebert, Verena Böhm, Julia K. Büttner, Rafaela Holtappels, Niels A. Lemmermann

**Formal analysis:** Stefan Ebert, Verena Böhm, Julia K. Büttner

**Funding acquisition:** Rafaela Holtappels, Matthias J. Reddehase, Niels A. Lemmermann

**Investigation:** Stefan Ebert, Verena Böhm, Julia K. Büttner

**Methodology:** Stefan Ebert, Verena Böhm, Wolfram Brune, Melanie M. Brinkmann

**Project administration:** Matthias J. Reddehase, Niels A. Lemmermann

**Supervision:** Rafaela Holtappels, Matthias J. Reddehase, Niels A. Lemmermann

**Validation:** Matthias J. Reddehase, Niels A. Lemmermann

**Visualization:** Stefan Ebert, Niels A. Lemmermann

**Writing – Original Draft Preparation:** Matthias J. Reddehase, Niels A. Lemmermann

**Writing – Review & editing:** Rafaela Holtappels, Wolfram Brune, Melanie M. Brinkmann

## Funding

This research was funded by the Deutsche Forschungsgemeinschaft, Collaborative Research Center (CRC) 490 (R.H. and M.J.R), Clinical Research Group KFO 296/2 (Project No. BR 1730/7-1; W.B.), and CRC 1292 (Project No. 318346496; M.J.R. and N.A.L.). N.A.L is a member of the DFG-funded Cluster of Excellence ImmunoSensation – EXC2151 – at the University of Bonn. MB is supported by the SMART BIOTECS alliance between the Technische Universität Braunschweig and the Leibniz Universität Hannover, an initiative supported by the Ministry of Science and Culture (MWK) of Lower Saxony, Germany, and the Helmholtz Association (W2/W3-090). The funders had no role in the design of the study, in the collection, analyses, or interpretation of data, in writing of the manuscript, or in the decision to publish the results.

## Data Availability Statement

All relevant data are within the manuscript and its Supporting Information files.

## Supporting information

**S1 Fig. Significant contribution of macrophages to the virus-specific CD8 T-cell response.** Viral epitope-specific CD8 T-cell response determined for the spleen on day 7 after intra-plantar infection of BALB/c mice (cohorts of 5 mice) with mCMV-WT. (Grey shaded bars) depletion of macrophages by clodronate liposomes, (open bars) control group left undepleted. For further information, see the legend of Fig 2B.

**S2 Fig. Time course of the viral epitope-specific CD8 T-cell response dependent on presence or absence of apoptosis.** Viral epitope-specific CD8 T-cell response determined daily for the RLN, specifically the popliteal lymph node, between day 2 and day 7 (d2-d7) after intra-plantar infection of BALB/c mice with mCMV-ΔM36 (ΔM36) lacking M36, and the revertant virus mCMV-ΔM36-Rev expressing M36. (Green color) apoptosis can take place. (Red color) apoptosis is blocked. For further information, see the legend of Fig 2B.

**S3 Fig. Release from necroptosis blockade by functional deletion of M45 leads to virus growth attenuation in the RLN but not locally at the viral entry site.** Levels of infection were determined by quantitation of IE1 transcripts in plantar (footpad) tissue (left panel) and in the draining RLN, the popliteal lymph node (right panel), at 48 hours (day 2) after intra-plantar infection (day 0) with the viruses indicated in the internal legend. For further information and statistical evaluation, see the legend of Fig 3.

## Notes

### Competing Interest Statement

The authors have declared no competing interest.

